# Presynaptic boutons that contain mitochondria are more stable

**DOI:** 10.1101/580530

**Authors:** Robert M. Lees, James D. Johnson, Michael C. Ashby

## Abstract

Addition and removal of presynaptic terminals reconfigures neuronal circuits of the mammalian neocortex, but little is known about how this presynaptic structural plasticity is controlled. Since mitochondria can regulate presynaptic function, we investigated whether the presence of axonal mitochondria relates to structural plasticity of presynaptic boutons in mouse neocortex. We found that the overall density of axonal mitochondria did not appear to influence loss and gain of boutons. However, positioning of mitochondria at individual presynaptic sites did relate to increased stability of those boutons. In line with this, synaptic localisation of mitochondria increased as boutons aged and showed differing patterns of localisation at *en passant* and *terminaux* boutons. These results suggest that mitochondria accumulate locally at boutons over time to increase bouton stability.

## Introduction

Individual cortical presynaptic terminals can be added and removed on axonal branches on a timescale ranging from days to years (De Paola et al., 2006; Grillo et al., 2013; Mostany et al., 2013; Qiao et al., 2015). Alterations in this presynaptic turnover are related to learning (Holtmaat and Caroni, 2016; Johnson et al., 2016; Ash et al., 2018) and disease (Jackson et al., 2017), showing their importance for plasticity of neural circuits. However, little is known about cellular control of bouton structural plasticity, although it has been suggested that mitochondria may play a role (Smit-Rigter et al., 2016).

Mitochondria and synaptic efficacy are strongly linked. Ultrastructural features of efficacy (e.g. postsynaptic density size or number of docked vesicles) are positively correlated to presynaptic mitochondria (Smith et al., 2016; Cserép et al., 2018; Kasthuri et al., 2015), mitochondrially-derived ATP is required to sustain neurotransmission during elevated levels of stimulation (Sobieski et al., 2017; Rangaraju et al., 2014; Hall et al., 2012) and mitochondria modulate presynaptic release by sequestering cytosolic calcium or altering ATP concentrations (Lewis et al., 2018; Vaccaro et al., 2017; Kwon et al., 2016; Sun et al., 2013). However, mitochondria only localise to a subpopulation of boutons (Smit-Rigter et al., 2016; Vaccaro et al., 2017; Obashi and Okabe, 2013; Kang et al., 2008; Chang et al., 2006) and very little is known about whether the spatial distribution of mitochondria relative to presynaptic sites is related to bouton formation, longevity or removal (Smit-Rigter et al., 2016). To address this, we have used chronic, *in vivo* two-photon (2P) imaging to investigate the relationship between mitochondrial localisation in axons and the structural plasticity of presynaptic boutons.

## Results

To monitor presynaptic bouton structure alongside axonal mitochondria, we transduced neurons of the mouse motor cortex with an adeno-associated virus (AAV) that co-expressed cytosolic EGFP and mitochondrially-targeted (MTS)-TagRFP (Figure 1A).

**Figure 1.**
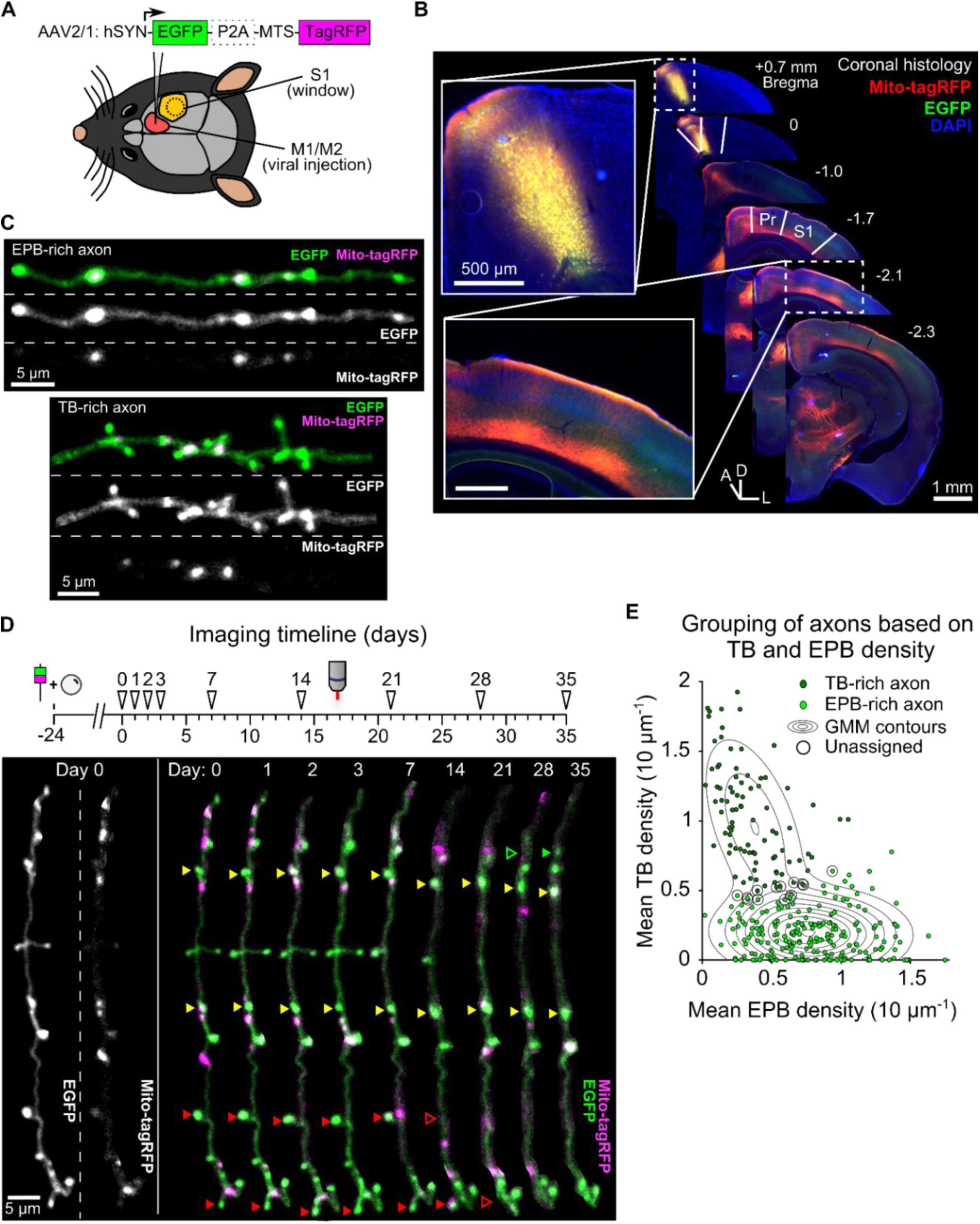
Tracking bouton plasticity and mitochondrial positioning in axons of motor cortex neurons. (**A**) AAV expressing cytosolic EGFP and a mitochondrial targeting sequence (MTS) conjugated to TagRFP was injected in to M1/M2 and a glass cranial window was implanted over S1. (**B**) A series of coronal brain slices showing viral injection site across the M1/M2 border (*inset, top*) and the axonal projection site at S1 under the cranial window (*inset, bottom*). Only the ipsilateral half of brain sections are shown. Pr = parietal cortex, D = dorsal, L = lateral, A = anterior. (**C**) Cropped 2P images from *in vivo* imaging show axons with high EPB (EPB-rich) or TB (TB-rich) densities. (**D**) (*top*) Imaging timeline for tracking bouton structural plasticity (loss and gain). Viral injection and cranial window implantation were performed 24 days prior to initial 2P imaging. Arrowheads indicate imaging timepoints. (*bottom*) Structure and mitochondrial localisation in a single cropped axon over 35 days imaged using *in vivo* 2P microscopy. Some boutons are labelled with arrowheads to show examples of stable (yellow), lost (red) or gained (green) boutons. (**E**) Gaussian mixture modelling (GMM) was used to determine two potential populations (*EPB-rich* and *TB-rich*) that result in the observed sample distribution of EPB and TB densities for each axonal segment (mean across time). Axons that had posterior probabilities below 70% were not assigned to a group (*circled*; see *Methods*). Contour lines indicate the slope of the GMM distribution.

A substantial projection from the motor cortex is made up of long-range axons that ramify in ipsilateral somatosensory cortex (Figure 1B; Veinante and Deschênes, 2003; Petreanu et al., 2009; Mao et al., 2011; Oswald et al., 2013). Imaging of intact, cleared brains showed that these axons travel over distances of more than 3 mm via cortical layers 5/6 or, to a lesser extent, superficially within layer 1 (Figure S1, S2). By placing a cranial window over somatosensory cortex, we imaged segments of these long-range axons within layer 1 using *in vivo* 2P microscopy. Mitochondria, putative *en passant* boutons (EPBs) and *terminaux* boutons (TBs) were clearly identified as increases in fluorescence intensity along the local axon backbone (Figure 1C, see Methods). We tracked structural synaptic plasticity of individual boutons by repeated imaging of the same axons at daily and weekly intervals over a total of up to 35 days (Figure 1D; n=12 animals, 306 axons). Visual inspection (Figure 1C) and statistical analysis based on the density of EPBs and TBs along each axon (Figure 1E) indicated that axons were mostly either EPB-rich or TB-rich. As this is the first characterisation of boutons in this axonal pathway, we compared the density and turnover of boutons in individual axonal branches (Figure S3). While there was a higher density of boutons in TB-rich axons, no differences were found in bouton turnover between EPB-rich and TB-rich axons (Figure S3).

As mitochondria can support presynaptic function, we assessed whether the numerical density of boutons and mitochondria are correlated in axons (population mean over time was 1.09 ± 0.41, 1 S.D., and 0.69 ± 0.23 per 10 µm, respectively; Figure 2A). For individual axonal segments (median length of 75 µm, Figure S4), there was a strong correlation between the densities of mitochondria and putative boutons (mean over time; Pearson’s correlation, R^2^ = 0.50, p = 1.16 × 10^-20^; Figure 2B). This suggested that the formation and/or elimination of boutons may relate to the mitochondrial population. To assess this, we compared the fraction of dynamic boutons (loss *and* gain) to mitochondrial density (Figure 2C). Bouton dynamics were not significantly correlated to mitochondrial density in axonal segments at daily or weekly intervals (R_s_ = 0.005, Spearman’s correlation, p = 0.944; Figure 2C, Table S1). Similarly, there was no apparent correlation between bouton dynamics and mitochondrion-to-bouton ratios (R_s_= 0.024, Spearman’s correlation, p = 0.733; Figure 2C, Table S1), indicating that the overall availability of mitochondria along a stretch of axon is not related to the degree of structural plasticity occurring there.

**Figure 2.**
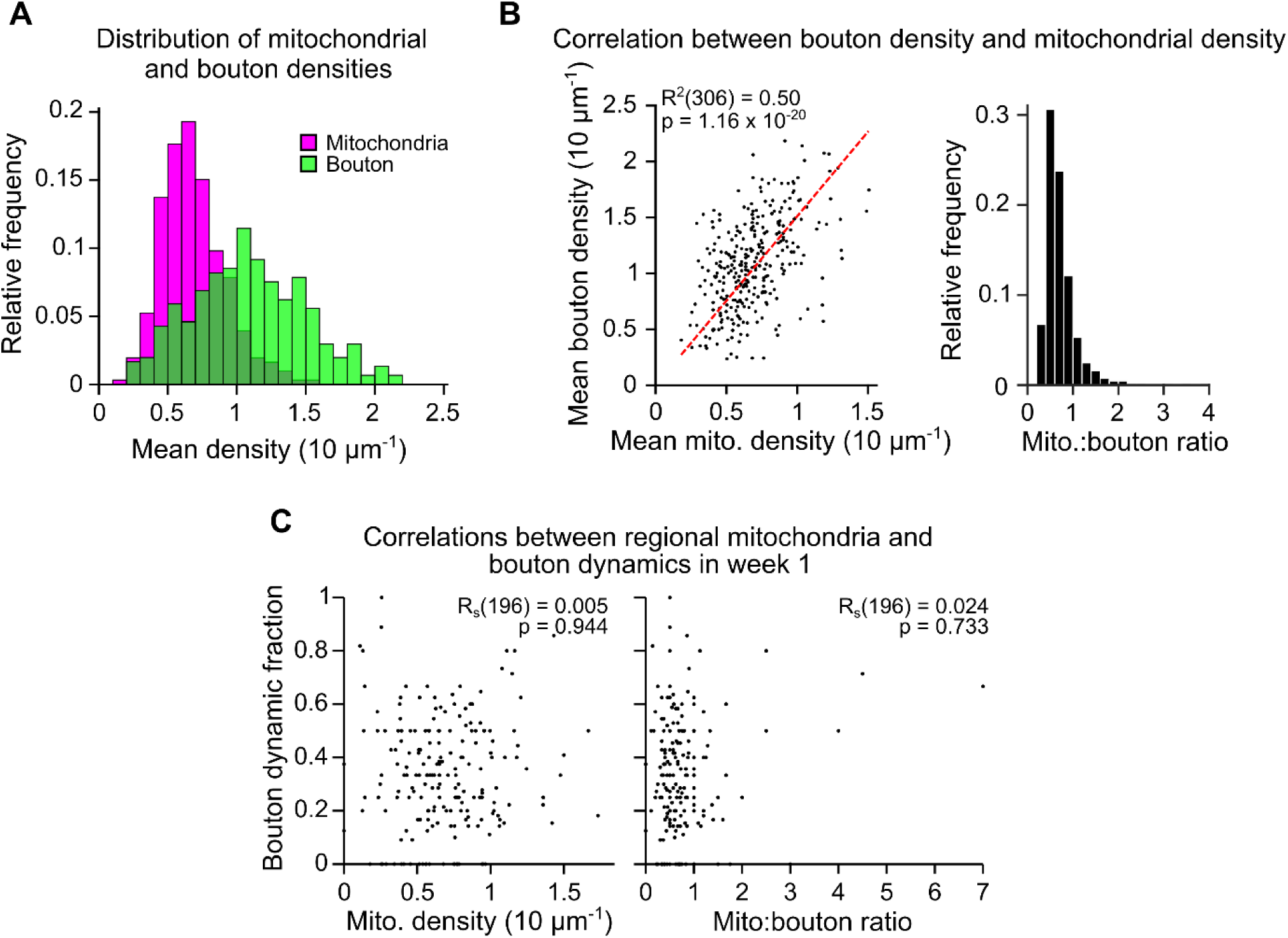
Mitochondrial density along an axonal segment is correlated to bouton density but not bouton dynamics. (**A**) Mitochondrial and bouton density distributions for all axons (mean across time, n = 306 axons). (**B**) (*left*) Bouton and mitochondrial densities for each axon were strongly correlated (mean across time; R^2^ = Pearson’s correlation, n = 306). *Red dashed line* = regression line. (*right*) The histogram shows the distribution of mitochondrion-to-bouton ratios for all axons with a median of 0.65 (approximately 2 mitochondria to every 3 boutons). (**C**) Example correlations between the fraction of boutons that were dynamic on each axon (lost or gained; *bouton dynamic fraction*) and either: (*left*) the number of mitochondria relative to the number of boutons (mitochondrion-to-bouton ratio), or (*right*) mitochondrial density. Results from the first weekly interval (between days 7 and 14) are shown (n = 196 axons over all weekly intervals, R_S_ = Spearman’s rank correlation, see Table S1).

As the overall density of axonal mitochondria was related to bouton density, but not to bouton dynamics, we assessed if there was instead a more local relationship between individual boutons and mitochondria near them. Based on previous studies and effective resolution limits of our imaging, we chose 1.5 µm as a biologically-relevant distance to presynaptic terminals (Smith et al., 2016; Smit-Rigter et al., 2016; see *Methods*). Whereas most mitochondria (65%) were found within 1.5 µm of presynaptic terminals (centroid-to-centroid distance, Figure 3A), only a minority of the total pool of putative boutons (44%) had mitochondria closer than 1.5 µm (Figure 3B). This local organisation did not occur by chance, as randomising or mirroring positions of either mitochondria or boutons along the axon backbone resulted in greater distances between them (Figure 3A, B, C). To determine if the structure of boutons affected the ability for mitochondria to localise there, we divided the bouton population in to EPBs and TBs.

**Figure 3.**
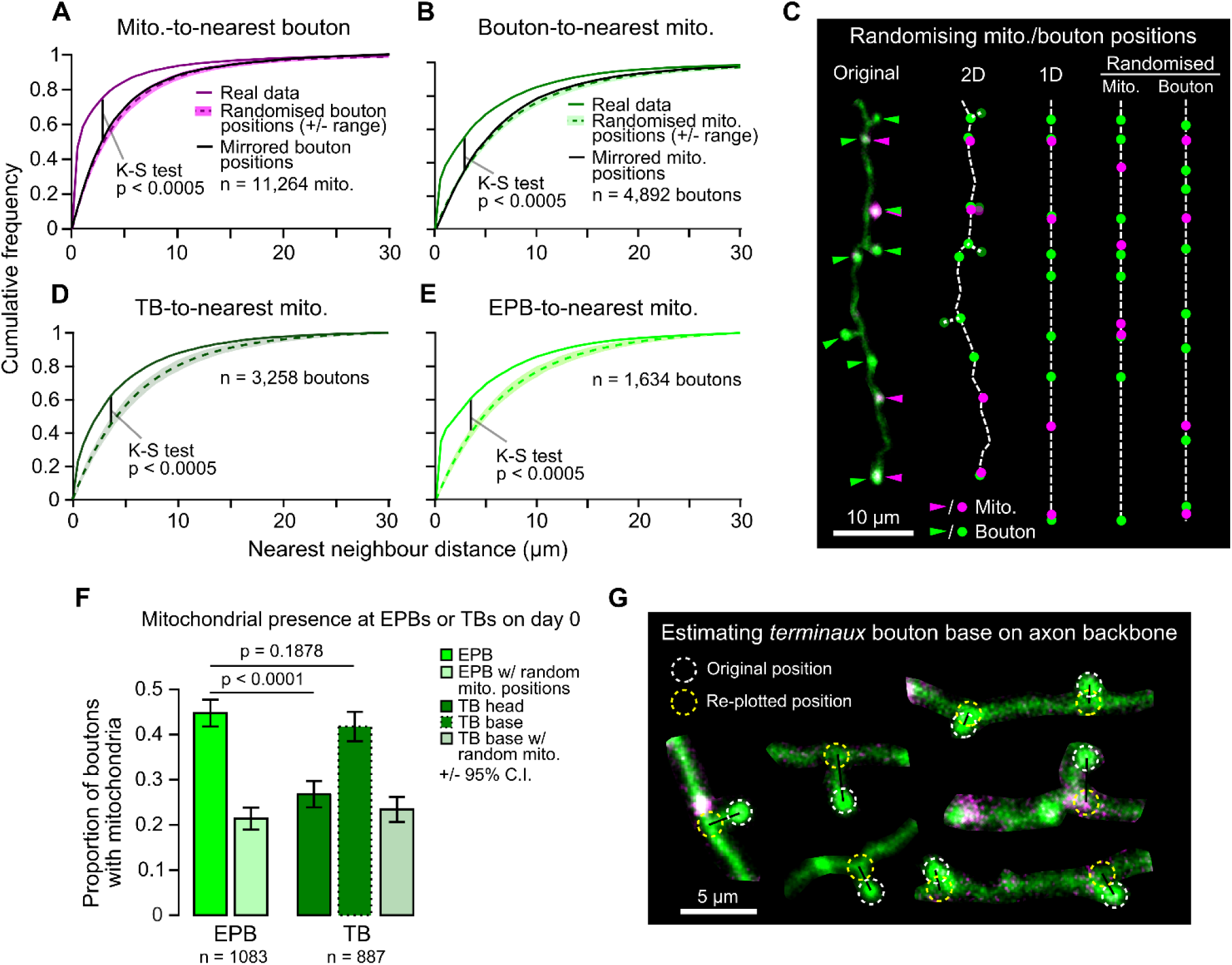
Mitochondria are positioned more closely to *en passant* boutons than *terminaux* boutons. (**A**) The distribution of distances between each mitochondrion and their nearest bouton was plotted against the results from 1000 rounds of randomised positioning of boutons for comparison to chance levels. Median ± range (*shaded area*). Kolmogorov-Smirnov (*K-S*) test between real data and the median of randomised positioning. As a further control, the real bouton positions were mirrored along the axon backbone to maintain the inter-bouton distances (*black line*) resulting in a similar distribution to the randomised positioning. (**B**) Same as in (A), but for boutons and their nearest mitochondrion compared to results from randomised/mirrored positioning of mitochondria. (**C**) Illustration of the routine for randomising positions. The original image was manually traced and a 2D skeleton interpolated from the segmented line trace. TBs were approximately placed at the nearest point on the axon backbone (their base) for randomising in 1D. The 2D skeleton was then straightened to 1D and either mitochondria were randomly positioned alongside real bouton positions, or *vice versa*. (**D-E**) Same as in (B), but for TBs only (D; using TB base position, see G) or EPBs only (E). (**F**) A greater proportion of EPBs have mitochondria within a biologically-relevant distance (1.5 µm, see *Methods*) than TBs (day 0 data; Chi-squared test). When mitochondrial localisation was considered from the TB base instead of the head the difference was lost (Chi-squared test). Error bars ± 95% C.I. (**G**) Estimated location of TB bases was achieved by finding the nearest neighbour point on the axon backbone that was closest to the TB head and re-plotting the TB to that position. 2P images were cropped for easier visualisation.

There was a higher likelihood of mitochondria at EPBs (45 ± 3%, 95% C.I.) than TBs (measured from TB head, 27 ± 3%; Chi-squared test, p < 0.0001; Figure 3F). It is possible that mitochondria reside near TBs, but do not traverse their neck region; therefore, we estimated the location of the base of TBs by re-plotting them to where they joined the axon backbone (Figure 3G). The probability of mitochondria at the base of TBs was higher (42 ± 3%) than at the head, and not different from that of EPBs (data from day 0; Chi-squared test, p = 0.188; Figure 3F, G).

Given that mitochondria have been implicated in the control of presynaptic function, we hypothesised that mitochondrial presence may relate to bouton maturity. To test this hypothesis, we separated boutons by age (*new* or *pre-existing*; Figure 4A). New boutons were those formed between daily imaging sessions (<24 hours old), whereas pre-existing boutons were present before imaging began (mixed ages). Some pre-existing boutons would have been formed in the previous day and should have been classed as new boutons, but we estimated this to be <10% of the total population as this was the rate of daily bouton formation (Figure S5). Pre-existing boutons were more likely than newly-formed boutons to have a resident mitochondrion (proportion with mitochondria on first day tracked, *pre-existing* 38 ± 2%, *new* 32 ± 4%, 95% C.I., Chi-squared test, p = 0.0024; Figure 4B). However, new boutons were still more likely to have mitochondria nearby than predicted by chance (17 ± 3%, 95% C.I., Chi-squared test, p < 0.0001), as were pre-existing boutons (17 ± 2%, Chi-squared test, p < 0.0001; Figure 4B). Further to this, the likelihood of mitochondrial presence at long-lived boutons (those that survived from the start) rose over time (*new* boutons 28 ± 12% to 43 ± 13%, *pre-existing* 46 ± 4% to 52 ± 4%, Cochran’s Q test, *new*, p = 0.235, *pre-existing*, p < 0.0005; Figure 4C). This was not the case with randomised mitochondrial positions, suggesting it is not a chance phenomenon (*pre-existing* 15 ± 3% to 18 ± 3%, *new* 13 ± 9% to 22 ± 11%, Cochran’s Q test, *new*, p = 0.578, *pre-existing*, p = 0.134). It was also not due to a general trend towards increased synaptic localisation of mitochondria over time, as this was stable across the imaging paradigm (Figure S6). These data show that the longer a bouton survives, the more likely it is to have a mitochondrion nearby.

**Figure 4.**
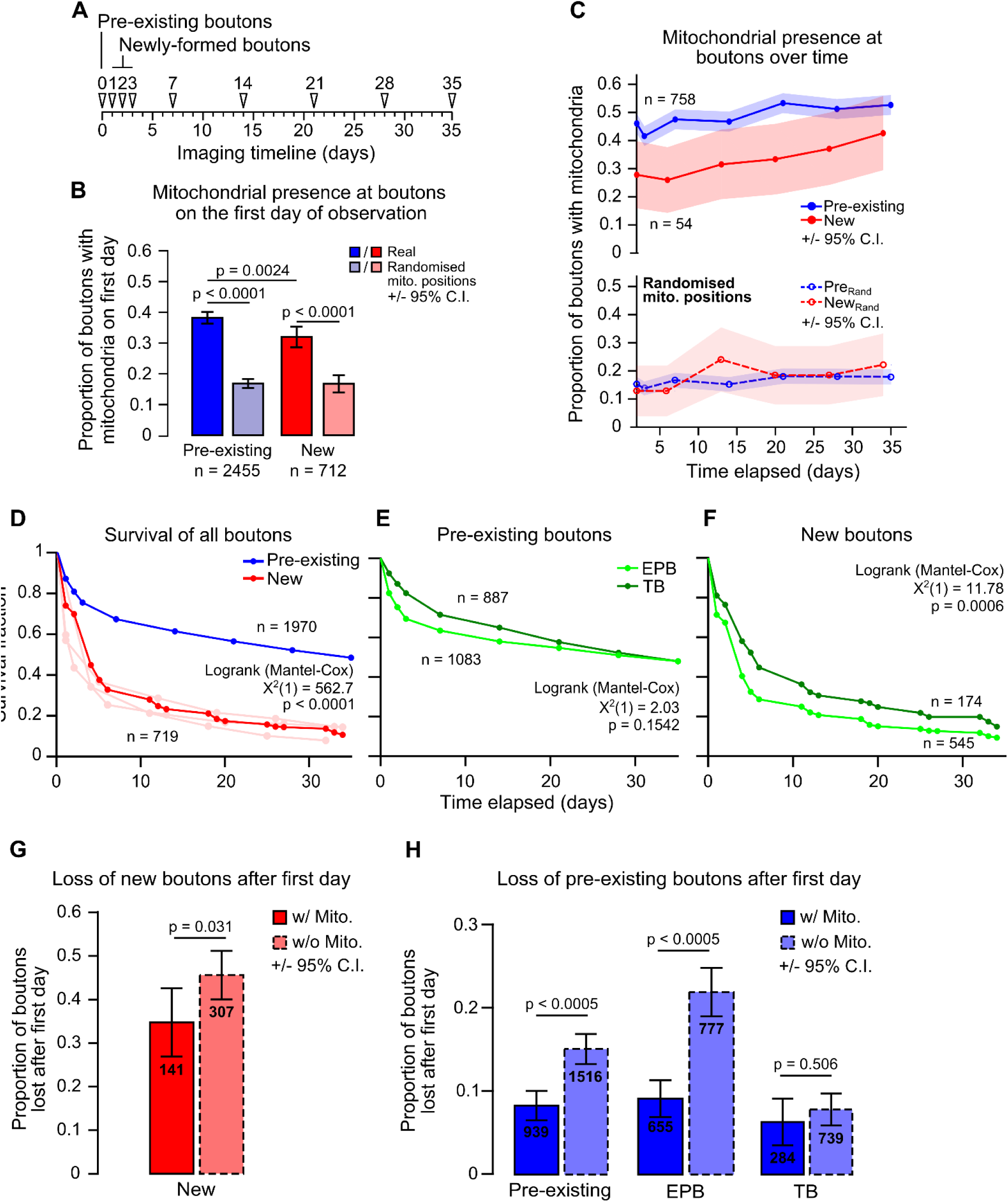
Mitochondrial presence at individual boutons is positively related to bouton age and longevity. (**A**) (*top*) Timeline indicating the classification of pre-existing boutons (first identified on day 0) and newly-formed boutons (first identified on days 1, 2 or 3). (**B**) Pre-existing boutons were more likely to have mitochondria (<1.5 μm) than newly-formed boutons (Chi-squared test). Newly-formed boutons had more mitochondria present than with randomised positioning of mitochondria, as did pre-existing boutons (Chi-squared test). Error bars ± 95% C.I. (**C**) Boutons that were present in every timepoint after day 2 (after all newly-formed boutons were identified) had their mitochondrial localisation tracked. Pre-existing boutons showed a significant increase in mitochondrial presence (Cochran’s Q test: X^2^(6) = 51.359, p < 0.0005). New boutons also showed an increase; however, this was not statistically significant (Cochran’s Q test: X^2^(5) = 6.81, p = 0.235). When mitochondrial positions were randomised, both new and pre-existing boutons did not show significant increases in mitochondrial localisation (Cochran’s Q test: New, X^2^(5) = 3.807, p = 0.578; pre-existing, X^2^(6) = 9.787, p = 0.134). Shaded areas are ± 95% C.I. (**D**) Survival of boutons was measured as the time until bouton loss. Pre-existing boutons were significantly more stable than new boutons (Logrank test). The new bouton population was pooled from three consecutive days (light red lines). (**E-F**) Of the pre-existing population, TB and EPB survival was similar (E), however there was a small significant decrease in survival for new EPBs compared to TBs (F; Logrank test). (**G**) The proportion of new boutons with or without mitochondria that were lost after their first day. There was a significant decrease in bouton loss when mitochondria were present at new boutons (Fisher’s exact test). (**H**) Similarly, pre-existing boutons with mitochondria were half as likely to be lost when compared to those without mitochondria (Fisher’s exact test). This relationship was due to the population of EPBs and not TBs. Error bars ± 95% C.I.

It has been shown that newly-formed cortical boutons tend to be lost more quickly than pre-existing boutons (Ash et al., 2018, Qiao et al., 2015). Here, we show this is also true for boutons on long-range axons of motor cortical neurons, where less than 30% of new boutons survived more than 1 week compared to ∼70% survival for pre-existing boutons (median survival: *new*, 4 days, *pre-existing*, 35 days, Logrank test, p < 0.0001; Figure 4D). Interestingly – despite their differing structures – TBs and EPBs show similar survival within these groups (Figure 4E, F), with only a small difference in median survival between new EPBs and TBs (*new EPB*, 4 days, *new TB*, 6 days, *pre-existing EPB* and *TB*, 35 days, Logrank test, *new*, p = 0.0006, *pre-existing*, p = 0.154; Figure 4F). This aligns with the similarity in overall bouton turnover rates between EPB-rich and TB-rich axons (Figure S3).

To determine if mitochondria relate to the stability of individual boutons locally, we assessed the survival of boutons with and without mitochondria. For new synaptic boutons, the chance of being removed was only slightly reduced if mitochondria were present (20% relative reduction after first day, *without mitochondria* 45 ± 6%, *with mitochondria* 36 ± 8%; Chi-squared, p = 0.031; Figure 4G). However, the effect was much more pronounced for older, pre-existing boutons, in which having mitochondria decreased the probability of subsequent removal by ∼60% (loss after first day, *without mitochondria* 17 ± 2%, *with mitochondria* 7 ± 2%; Chi-squared, p < 0.0005; Figure 4H). This relationship between mitochondrial proximity and enhanced bouton survival was consistent across time for pre-existing boutons (Figure S7). As localisation of mitochondria at TBs and EPBs appeared to be different, we assessed the impact of having resident mitochondria on the survival of the two bouton types separately for pre-existing boutons. Indeed, mitochondrial presence was strongly related to decreased loss of EPBs, but not of TBs (Figure 4H). These results suggest that immediate survival of new boutons is weakly related to local mitochondrial presence, but this relationship becomes stronger and more consistent as boutons age.

## Discussion

It has long been reported that many, but not all, presynaptic release sites have mitochondria in close proximity to them (Gray, 1959; Shepherd and Harris, 1998; Chang et al., 2006; Kang et al., 2008; Obashi and Okabe, 2013; Smit-Rigter et al., 2016; Vaccaro et al., 2017). We found that axonal mitochondria in motor-somatosensory neurons are also preferentially associated with a subpopulation of synaptic boutons (Figure 3, 4B-C), suggesting bouton-specific recruitment and/or anchoring mechanisms (Kang et al., 2008, Courchet et al., 2013). The fact that mitochondria can modulate synaptic function suggests that having a resident mitochondrion may also relate to the activity-dependent plasticity of the synapse. Here, we have shown that there is indeed an association between mitochondrial positioning at presynaptic terminals and their structural longevity. As with other axons (De Paola et al., 2006; Qiao et al., 2015; Ash et al. 2018; Morimoto et al., 2018), we found that these motor-somatosensory axons exhibit structural plasticity driven by turnover of a minority of their synaptic boutons (Figure 4D-F). Newly-formed boutons are more likely to possess mitochondria within their first 24 hours (our smallest imaging interval) than by chance (Figure 4B), suggesting a link between synaptic and mitochondrial function even in the early stages of the synaptic lifecycle. Long-lasting boutons were even more likely to have resident mitochondria (Figure 4B, C) and this decreased their chance of being removed by half (Figure 4H). This results in a persisting population of synaptic boutons that contain mitochondria. However, having a mitochondrion is not absolutely required for bouton survival as not all boutons that survived across the 35 days of imaging had resident mitochondria. As such, it seems likely that mitochondrial recruitment links to some synaptic function that promotes synaptic longevity (Rangaraju et al., 2019). It remains unknown whether mitochondria directly influence long-term plasticity of synaptic function, as recently shown within dendrites (Divakaruni et al., 2018; Smith et al., 2016), or are simply recruited by alterations in synaptic activity to support presynaptic function (Vaccaro et al., 2017).

Our data suggest that any link between mitochondria and plasticity is local to neighbouring synapses. This is because, although the density of mitochondria along different axonal branches varied considerably, it did not correlate with rates of bouton plasticity at the branch level (Figure 2). In contrast, close proximity of mitochondria (within 1.5 μm) did relate to individual bouton stability (Figure 4). Furthermore, we found that mitochondria were generally located closer to EPBs than to TBs (Figure 3). This may be because physical access to the bouton head is restricted by the neck of TBs or could reflect functional differences between bouton types. Indeed, the difference in proximity was mirrored by the fact that local mitochondria were strongly linked to survival of EPBs but not TBs. Potentially divergent plasticity mechanisms at TBs and EPBs highlights the need for further investigation of the underexplored differences between different axonal bouton types.

## Acknowledgements

RML was funded by a Wellcome Trust PhD studentship (105386). JDJ was funded by a BBSRC-CASE PhD studentship (1370828). Equipment used in MCA’s laboratory was partially funded by the Medical Research Council (MR/J013188/1) and EUFP17 Marie Curie Actions (PCIG10-GA-2011-303680). We thank the Wolfson Bioimaging Facility for their support and expertise and MRC funding of a preclinical in vivo functional imaging platform for translational regenerative medicine. We thank Lilly UK for the gift of the AAV used in this study.

## Author contributions

RML, JDJ and MCA designed, developed and carried out the experiments and analysis. RML and MCA wrote the manuscript.

## Materials

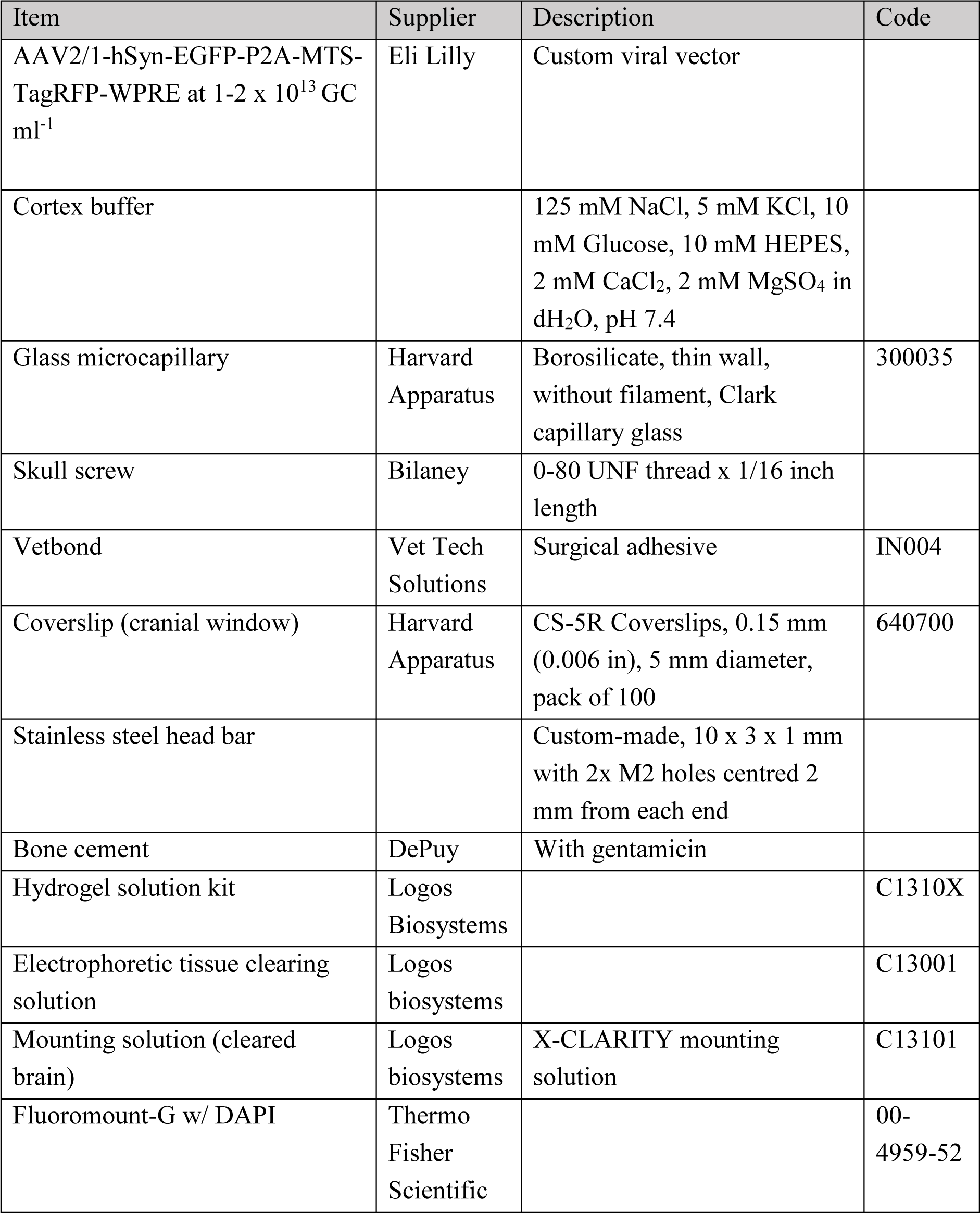

| Item | Supplier | Description | Code |
| --- | --- | --- | --- |
| AAV2/1-hSyn-EGFP-P2A-MTS-TagRFP-WPRE at $1-2 \times 10^{13}$ GC ml<sup>-1</sup> | Eli Lilly | Custom viral vector | |
| Cortex buffer |  | 125 mM NaCl, 5 mM KCl, 10 mM Glucose, 10 mM HEPES, 2 mM CaCl<sub>2</sub>, 2 mM MgSO<sub>4</sub> in dH<sub>2</sub>O, pH 7.4 |  |
| Glass microcapillary | Harvard Apparatus | Borosilicate, thin wall, without filament, Clark capillary glass | 300035 |
| Skull screw | Bilaney | 0-80 UNF thread x 1/16 inch length |  |
| Vetbond | Vet Tech Solutions | Surgical adhesive | IN004 |
| Coverslip (cranial window) | Harvard Apparatus | CS-5R Coverslips, 0.15 mm (0.006 in), 5 mm diameter, pack of 100 | 640700 |
| Stainless steel head bar |  | Custom-made, 10 x 3 x 1 mm with 2x M2 holes centred 2 mm from each end |  |
| Bone cement | DePuy | With gentamicin |  |
| Hydrogel solution kit | Logos Biosystems |  | C1310X |
| Electrophoretic tissue clearing solution | Logos biosystems |  | C13001 |
| Mounting solution (cleared brain) | Logos biosystems | X-CLARITY mounting solution | C13101 |
| Fluoromount-G w/ DAPI | Thermo Fisher Scientific |  | 00-4959-52 |

## Equipment

| Item | Supplier | Description |
| --- | --- | --- |
| Two-photon microscope (*in vivo* imaging) | Scientifica | Custom-built with filter cube carousel and epifluorescence capabilities |
| Ti-Sapphire laser | Spectra-Physics | MaiTai Ti-Sapphire tuneable laser (680-1040 nm) |
| 60x water-immersion objective | Olympus | For *in vivo* 2P imaging, 0.9 NA |
| X-CLARITY tissue clearing system | Logos Biosystems |  |
| Two-photon microscope (cleared brain imaging) | Leica | Leica SP8 AOBS laser-scanning microscope w/ Leica DM6000 upright epifluorescence microscope |
| Ti-Sapphire laser | Spectra-Physics | MaiTai DeepSee Ti-Sapphire tuneable laser (680-1300 nm) |

## Software

| Software | Developer | Version | Description |
| --- | --- | --- | --- |
| ScanImage | Vidrio Technologies | 5.1 | Two-photon microscope control |
| Micromanager | Open Imaging | 1.4 | Camera control |
| ImageJ (FIJI package) | ImageJ Development Team | 1.51a (2.0.0-rc-43) | Image processing and analysis |
| MATLAB | MathWorks | R2016a | Data processing and analysis |
| Prism | GraphPad | 7 | Graph making and basic statistical testing |
| G* Power | Universität Düsseldorf |  | Statistical power calculations |
| SPSS Statistics | IBM | 24 | Statistical testing |

## Methods

### Animal husbandry

All procedures involving animals were carried out adhering to the Animals (Scientific Procedures) Act 1986 as outlined by the Home Office, UK and approved by the University of Bristol Animal Welfare and Ethics Review Board.

Adult (2.5 months old) C57Bl/6 male mice were used for all experiments, living on a 12-hour light-dark cycle. Animals were housed individually to avoid loss of the cranial window implant due to fighting. Large (∼30 x 50 x 25 cm) cages were used and extra enrichment was provided for each cage, consisting of tunnels, shelters, wheels and foraging food to increase experience-dependent turnover of presynaptic terminals (Landers et al., 2011, Nithianantharajah et al., 2004, Briones et al., 2004).

### Viral DNA construct

The virus used for intracranial injection was a custom-made adeno-associated virus of serotype 2/1 expressing a bi-cistronic vector (AAV2/1-hSYN-EGFP-P2A-MTS-TagRFP). The human synapsin promoter (hSYN) was used to limit expression to neuronal cells. Cytosolic enhanced green fluorescent protein (EGFP) was separated by a P2A peptide from mitochondrially-targeted tagRFP (red fluorescent protein, fused to amino acids 1-29 of Cox8a subunit of cytochrome oxidase), which localised to the inner mitochondrial membrane. The P2A peptide is a self-cleaving peptide of the 2A family from porcine teschovirus, that has a high cleaving efficiency (Kim et al., 2011). Additionally, Woodchuck Hepatitis Virus Posttranscriptional Regulatory Element was used to increase protein expression (Zuffery et al., 1999). The viral titre used for injections was in the range of 1-2 x 10^13^ particles mL^-1^ in cortex buffer (125 mM NaCl, 5 mM KCl, 10 mM Glucose, 10 mM HEPES, 2 mM CaCl_2_, 2 mM MgSO_4_ in dH_2_O, pH 7.4).

### Surgery

To reduce stress, animals were allowed at least one week to acclimatise to unfamiliar environments after relocation before the commencement of procedures. Intraperitoneal injections of rimadyl (analgesic, 4 mg mL^-1^ kg^-1^) and dexamethasone (anti-inflammatory, 0.5 mg mL^-1^ kg^-1^) were given pre-operatively to reduce pain and inflammation. Aseptic technique was used to limit the possibility of infection, guidelines were followed as outlined by the Laboratory Animal Science Association (http://www.lasa.co.uk/wp-content/uploads/2017/04/Aseptic-surgery-final.pdf). The protocol described in Holtmaat et al. (2009) was followed for cranial window implantation, which is briefly described below with amendments.

Animals were anaesthetised using gaseous isoflurane at 3-4% for induction and 1-2% to sustain anaesthesia throughout surgery, carried by O_2_. The top of the head was shaved and placed in a stereotaxic frame, and the scalp and periosteum were removed. The skull bone was kept moist throughout surgery with cortex buffer.

The intracranial viral injection site was measured +0.7 mm lateral (always to the right) and +1.0 mm anterior from Bregma, as these coordinates corresponded to the primary/secondary motor cortex (Lein et al., 2007; Petreanu et al., 2012). A small (∼0.5 mm diameter) burr hole was made in the skull using a high-speed motorised hand drill. For viral injections, a glass capillary tube was pulled in to a micropipette with a long, pointed tip and bevelled on a whetstone to sharpen it further. Virus was injected intracranially using a Hamilton syringe and motorised pump at a rate of 100 nL/min. A volume of 300 nL was injected at depths of 300 μm (first) and 700 μm (second) from the pial surface to spread it across all cortical layers. Virus was allowed to spread for 3 min before moving the micropipette.

After viral injection, a screw (0-80 UNF thread, 1/16 inch length) was implanted in the left parietal skull bone to anchor the cranial window implant to the skull. Subsequently, a thin layer of VetBond™ glue was spread across the skull, to the skin edges, avoiding the right parietal skull bone where the cranial window was to be implanted. A 3-4 mm diameter craniectomy was made using a motorised hand drill centred on +2.5 mm lateral and −1.8 mm anterior of Bregma. A 5 mm circular glass coverslip was then secured over the craniectomy on top of a small volume of cortex buffer using VetBond™ glue.

Quick-drying bone cement (with gentamycin, DePuy) was used to apply a 1-2 mm thick layer of cement over the layer of VetBond™ glue. Cement was spread just over the edge of the coverslip as well as up to the edges of the skin incision. A stainless-steel bar (10 x 3 x 1 mm; used for securing the head during *in vivo* imaging) was placed over the left hemisphere as close to the cranial window as possible while leaving enough space for microscope objective access.

Animals were left for ∼24 days before imaging to allow for any inflammation to clear under the window and to allow viral expression.

### *In vivo* imaging

For *in vivo* imaging, a customised Scientifica upright two-photon microscope was used along with a motorised stage to aid precise movement in coordinate space for relocation of regions of interest (ROIs). Epifluorescence was used for low resolution mapping of expression across the window to guide two-photon imaging. Two MaiTai Ti-Sapphire tuneable lasers (tuneable from 680–1040 nm, Spectra Physics) were used and attenuation of laser power was controlled through either a Pockel’s cell or half-wave plate. Two-photon excitation wavelengths for imaging were typically 920 nm (EGFP) and 1040 nm (TagRFP). Laser lines were combined using a polarising beamsplitter cube in reverse, and combined power never exceeded 60 mW at the back focal plane of the objective. Acquisition was controlled by ScanImage software (Pologruto et al., 2003, version 5.1) and Micromanager software (Edelstein et al., 2014, version 1.4). Objective lenses used: 4x air 0.15 NA, 10x water-immersion 0.6 NA and 60x water-immersion 1.1 NA. Emission filter sets used for PMTs were BP 620/60 nm for TagRFP and BP 525/50 nm for EGFP. The stage was fitted with a micromanipulator for precise head fixation and rotation in every repeated imaging session using the implanted steel bar on the animal’s head, increasing ROI relocation efficiency. During *in vivo* imaging, mice were anaesthetised by gaseous isoflurane anaesthetic (1–2% carried by O_2_) and breathing was monitored to judge depth of anaesthesia. Breathing was kept in the range of 80-100 beats per minute by eye.

For each mouse, a large blood vessel bifurcation was chosen using reflected light and set as the origin for recording coordinates of ROIs. Two-photon imaging was used to locate ROIs based on the following criteria: sparse labelling, to reduce background and contamination from crossing axons; distinctive axonal structures, for easy relocation; distance from other ROIs, to increase the diversity of sampling.

Z-stacks of 20–50 μm were acquired at each ROI with a step size of 1 μm (60x, 1.1 NA objective). Images were acquired with 3x frame averaging, 1 us pixel dwell time at 1024 x 1024 pixels and a field of view of 76 x 76 μm, resulting in a final pixel size of 74 nm. Signal was matched between sessions by adjusting laser power because of differences in window quality between imaging sessions, which altered the signal-to-noise ratio. Up to 7 ROIs were chosen per animal and each imaging session was kept between one and two hours.

Axons were tracked for up to 35 days after the initial session, for a total of nine sessions (ethical limit) or until the cranial window was no longer optically clear due to bone regrowth or dural thickening. A small proportion of ROIs were first tracked at days one and two, rather than day zero of the imaging paradigm. Most ROIs and animals were tracked for the entire imaging time series.

### Histology

Following the end of an imaging paradigm, mice were administered a dose of 70– 100 μL of Euthatal (200 mg/ml sodium pentobarbital) intraperitoneally to achieve terminal anaesthesia. When the animal was deeply anaesthetised, exsanguination was performed, and the animal was transcardially perfused with 5-10 mL of 0.01 M PBS. This was followed by infusion of 20–30 mL of 4% paraformaldehyde (PFA) in 0.01 M PBS. The brain was then dissected out and post-fixed in 4% PFA in 0.01 M PBS at 4°C.

### Tissue clearing

Tissue clearing was carried out by following the protocol described in Lee et al. (2016), which is briefly outlined below. The brain was post-fixed for 24 hours in 4% PFA followed by overnight incubation in hydrogel solution (4% w/v acrylamide without bis-acrylamide, 1% w/v VA-044 initiator in 0.01 M PBS) at 4°C. Oxygen was removed from the solution by degasification using pure nitrogen bubbling through the solution (providing some agitation). Polymerisation of the hydrogel was carried out at 37°C in a water bath for ∼3 h. The brain was then mounted inside the X-CLARITY electrophoresis chamber (Logos Biosystems) in electrophoretic tissue clearing solution (4% SDS and 200 mM boric acid). The X-CLARITY machine was used according to the manufacturer’s instructions.

### Tissue sectioning

Histological sectioning was achieved using either a vibratome or freezing microtome. For vibratome sectioning, brains were embedded in 2% agarose (in distilled H_2_O), trimmed to the region of interest and series of 50 μm-thick sections were cut in 0.01 M PBS on a vibratome. For freezing microtomy, brains were incubated in a 30% sucrose solution (w/v) for up to 1 week. The brains were then sectioned in optimal cutting temperature (OCT) solution. Sections were directly mounted on glass microscope slides with No. 1.5 coverslips using Fluoromount-G containing DAPI nuclear stain.

### Imaging of tissue sections

Imaging of whole tissue sections was carried out on a widefield microscope (Leica DMI6000) with a mercury lamp and CCD camera (Leica DFC365FX monochrome) using Leica LAS X software. Filter sets were assigned for the following fluorophores: DAPI (Ex. 350/50 nm, 400 nm dichroic mirror, Em. BP 460/50 nm), EGFP (Ex.: 480/40 nm, 505 nm dichroic mirror, Em. BP 527/30 nm), TagRFP (Ex. 620/60, 660 nm dichroic mirror, Em. BP 700/38). Objective lenses used: 5x dry 0.15 numerical aperture (NA) and 20x dry 0.4 NA. Brightfield and DAPI signal of coronal or sagittal tissue sections were compared to the Paxinos Mouse Brain Atlas (Franklin et al. 2008) or Allen Mouse Brain Atlas (Lein et al., 2007) as a reference to confirm the positions of viral injections and window sites.

### Imaging of cleared tissue

The cleared brain was immersed in a small volume (5–10 mL) of mounting medium (X-CLARITY mounting solution) inside a 50 mL Falcon tube for at least 2 h before mounting. It was then placed in the centre of a circular wall of blu-tac inside the lid of a 35 mm dish to create a water-tight well. The well was filled partially with fresh X-CLARITY mounting medium and the chamber was sealed on top with a 35 mm coverslip pressed in to the blu-tac. The chamber was filled from a small inlet in the blu-tac using a 200 μL pipette and the inlet was sealed by squeezing the blu-tac back together.

Cleared tissue was imaged using a Leica SP8 AOBS confocal laser scanning microscope attached to a Leica DM6000 upright epifluorescence microscope with a Ti-Sapphire laser (MaiTai DeepSee; tuneable from 680-1300 nm) and a fixed-wavelength 1040 nm laser. Two hybrid GaAsP detectors were used with a BP 525/50 nm filter for EGFP and BP 630/75 nm for TagRFP. Objectives lenses used: 10x water-immersion 0.3 NA and 25x water-immersion 0.95 NA. Large z-stack mosaic images (5 μm steps for ∼1 mm) were acquired using the tilescan function in Leica LAS X software. The laser intensity was attenuated at shallower imaging depths to maintain signal-to-noise ratio. The images were then resliced to obtain the correct viewing angle.

### Image processing

*In vivo* images were processed using the FIJI package for ImageJ (Schindelin et al., 2012, version 2.0.0-rc-43/1.51a) and a custom ImageJ macro. The macro allowed for automated processing of images for each ROI, carrying out the following functions: (1) alignment of the EGFP signal within a single z-stack to correct drift and application of the transformation to the TagRFP channel using the MultiStackReg registration plugin, (2) matching of z-stack sizes between timepoints by addition of blank slices, (3) alignment of z-stacks between timepoints in the x-and y axes using maximum z-projections and the MultiStackReg plugin, (4) alignment of z-stacks in the z-axis using an edited version of the Correct 3D Drift plugin to only include the z-axis transformation. This resulted in a 5-dimensional (5D - XYZCT) stack of each ROI aligned to within 5 μm in x, y and z for both channels across all timepoints. For presentation in figures only, images were cropped and had brightness and contrast adjusted and a median filter (74 nm kernel) applied.

### Data quantification

Axonal segments were manually traced using the segmented line tool with spline as part of a custom ImageJ macro script. After tracing at each timepoint, a minimum volume that encompassed the axon across all the timepoints was cropped from the original 5D stack. Between 1 and 12 axonal segments were chosen from each ROI. Factors used to choose axonal segments were: good signal-to-noise (subjective measure by analyst), few crossing axons and existence in all timepoints.

Identification and indexing of presynaptic terminals and mitochondria were carried out manually on each cropped axonal segment using a custom ImageJ macro script and the multi-point tool. A gaussian blur (sigma = 2 pixels, 154 nm) was used to smooth the signal and presynaptic terminals were scored subjectively, using the local intensity profile as a guide (further information below). The position of each object (bouton or mitochondrion) was estimated from a point placed by the analyst.

Boutons were tracked across imaging sessions from the first timepoint they were identified. Boutons in separate timepoints were linked if they were in the same place relative to fiducial markers, including any crossing axons, kinked structure or other persistent boutons. Any bouton that was lost from the field of view for one timepoint (through a shift in alignment in the x-or y axes) was excluded entirely. All boutons were scored blind to the mitochondrial signal.

*En passant* boutons (EPBs) were larger in volume than the axon backbone and therefore had higher intensity relative to the backbone due to increased numbers of fluorescent molecules (cytosolic EGFP). An EPB had to have contiguous pixels in the x-, y-and z-axes to ensure it was not the result of noise. The intensity profile of an EPB needed to include sharp edges (relatively steep curve either side of peak) to exclude gentle changes in the axonal thickness. If the peak of an EPB was twice that of the local axon backbone (1.5 μm either side of the bouton) at any timepoint, the bouton was scored as being present. A bouton was scored as lost if it was below 1.3 times the local axon backbone. These criteria have been shown to be faithful indicators of synapse presence in correlative light and electron microscopy studies (Grillo et al., 2013; Song et al., 2016).

*Terminaux* boutons were scored as unilateral protrusions from the axon backbone with a bulbous appearance and sometimes consisted of a resolved thin neck that extended for less than 5 μm. Those extending for longer than 5 μm were considered to be axonal branches (Grillo et al., 2013).

A small proportion of boutons changed bouton type (*en passant* or *terminaux*) over the time series and so those boutons were classified based on their predominant type.

Mitochondria were identified as discrete objects that were 2x the global median background signal, with contiguous pixels in the x-, y-and z-axes and steep edges to their intensity profile. The axonal EGFP signal was used to verify that each mitochondrion was inside the axon only after it was scored.

### Data analysis

*A priori* power calculations were performed in G*Power software (Faul et al., 2007) to calculate the number of newly-formed boutons required to detect a 10% difference in survival between the two mitochondrial conditions (less than or greater than 1.5 μm from a bouton). This calculation resulted in an estimated sample size of 450 newly-formed boutons. The number of animals required to achieve this was estimated from pilot studies to be 10-15 animals. In this study, 21 animals were used, 15 were imaged and 12 produced high-quality data that was included (see exclusion criteria below).

The final dataset was obtained from three different batches of littermates. A total of 51 ROIs and 306 axons were tracked. The total number of mitochondria counted across all timepoints was 11,264 along with 4,892 unique boutons. Mitochondria were not linked between timepoints because they lacked individuality due to their ability to move, fuse and split.

Some data were excluded from the final dataset. Any ROI that was too dim for accurate axon tracing (subjectively based on analyst experience) within the first four timepoints was not tracked. Any axon where the signal-to-noise in a session was low enough that the scorer could not be confident in bouton scoring was removed. Data from one animal that had only 2 axons tracked was also removed.

The bouton dynamic fraction was calculated as the proportion of unique boutons on an axon that were either lost or gained. Specifically, the sum of gains and losses divided by the total number of unique boutons across the two timepoints: (gained + lost)/(gained + total_time1_).

### Mitochondrion and bouton co-localisation

Mitochondria were classified as being present at a bouton if the distance between their centroids (defined by points placed by the analyst) was less than or equal to 1.5 μm. A dichotomous variable (mitochondria present or not) was chosen for analyses rather than a continuous variable (distance from nearest mitochondrion) because the axonal segment was a small sample of the axonal arbor and the true distance to the nearest mitochondria from each bouton could not be accurately measured, especially for boutons at the edge of the field of view. The distance of 1.5 μm was biologically-relevant because a distance-dependent relationship with synaptic ultrastructure has been seen up to 3 μm away (edge of vesicle pool to edge of mitochondrion) using electron microscopy (Smith et al., 2016). Stronger effects on synaptic ultrastructure were seen with closer distances of mitochondria. The accuracy of the measured distance in our study was limited due to the resolution of light microscopy and accuracy of point placement by the analyst, therefore only one distance was chosen in the middle of the range (0-3 μm).

Randomisation of bouton or mitochondrion position was carried out in a similar fashion to Smit-Rigter et al. (2016). Axons were first plotted in two dimensions in MATLAB using interpolation from segmented line coordinates recorded in ImageJ. The length of the axon was then estimated using Euclidean distances and a line was created and split in to segments of 74 nm (the original pixel size of the 2P images). The real positions of the objects of interest (mitochondria or boutons) were plotted to the closest segment of the axon based on where they were in the original image using nearest neighbour distance calculations. Either mitochondria or boutons were then removed and randomly re-plotted along the axon without being placed closer than 1 μm together. Intervals of at least 1 μm were chosen to attempt to match the resolution limit with which two objects could be resolved using the 2P microscope in this study. This was repeated 1000 times for each axon and the range plotted.

### Statistics

Statistics were calculated using MATLAB (release 2016a), GraphPad Prism 7 or SPSS (IBM). Statistical significance was set at p < 0.05. Confidence intervals for proportions were calculated using the formula for single samples. The z value for a 95% coverage of a gaussian distribution is 1.96. Therefore, the formula is as follows: prop1 ± z*sqrt(prop1*(1-prop1)/N), where prop1 is the proportion, z is 1.96 and N is the number of samples in the population. Errors are given as 95% confidence intervals for proportions and standard deviation for all other data, unless otherwise stated.

For repeated measures statistical tests, group sizes were matched by only including axons present in all relevant timepoints for the particular test. To avoid pseudo-replication, samples were not pooled together across time from repeated measures.

A Gaussian mixture model was used to calculate posterior probabilities of axons being in EPB-rich or TB-rich groups based on EPB and TB densities. An assumed number of two gaussian components was defined by the analyst. Axons that fell under the threshold of 0.7 probability for both groups were not assigned a group.

Kaplan-Meier curves were created for survival analysis, based on time-to-event data. For bouton survival, this was the time from first observation until the bouton was no longer observed. Boutons that were no longer observed due to reasons other than loss were classed as “censored”.

## SUPPLEMENTARY MATERIAL

### Supplementary Table

**Table S1.** Spearman’s rank correlation at each timepoint to assess the correlation between mitochondrial density and the fraction of dynamic boutons. Weak and non-significant correlations between bouton dynamic fraction and mitochondrion-to-bouton ratio or mitochondrial density were seen. R_S_ = Spearman’s rank correlation, n = 196 axons over all weekly intervals, n = 224 over all daily intervals.

| Mitochondrion-to-bouton ratio |  |  |  | Mitochondrial density |  |  |  |
| --- | --- | --- | --- | --- | --- | --- | --- |
| Daily |  | Weekly |  | Daily |  | Weekly |  |
| $R_s$ | p-value | $R_s$ | p-value | $R_s$ | p-value | $R_s$ | p-value |
| 0.101 | 0.133 | 0.024 | 0.733 | 0.012 | 0.857 | 0.005 | 0.944 |
| 0.123 | 0.066 | 0.033 | 0.646 | -0.013 | 0.843 | -0.079 | 0.273 |
| 0.063 | 0.348 | 0.100 | 0.164 | -0.050 | 0.457 | -0.020 | 0.779 |
|  |  | 0.066 | 0.357 |  |  | -0.100 | 0.164 |

### Supplementary Figures

**Figure S1.**
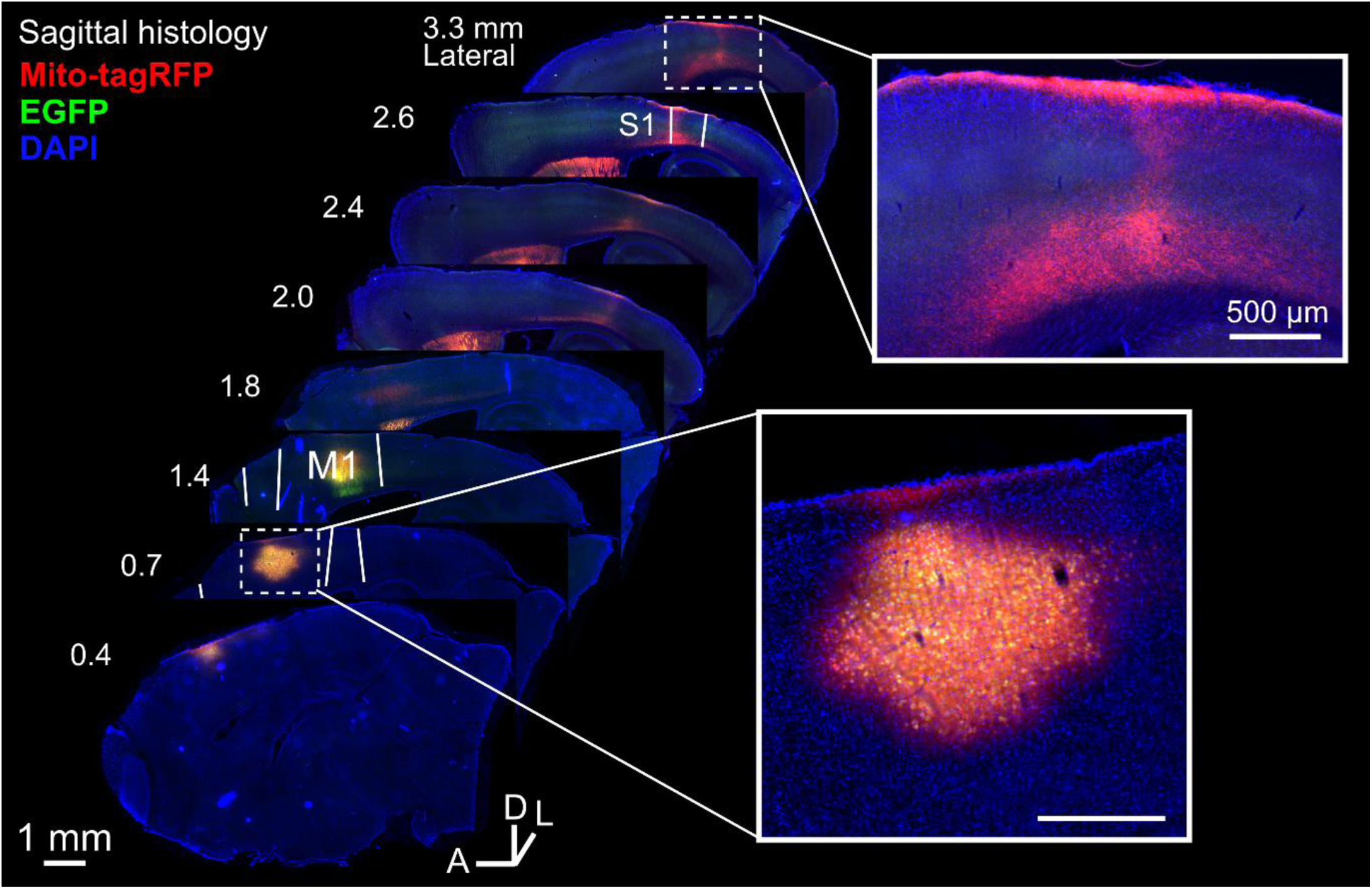
Ipsilateral motor-somatosensory axonal projections. Widefield images of a sagittal series of histological brain slices shows the viral injection site (*bottom inset*) between primary and secondary motor cortex (M1 and M2). Labelled cells projected their axons to subcortical areas as well as through deep cortical layers and ultimately arborising in superficial layers of somatosensory cortex (S1).

**Figure S2.**
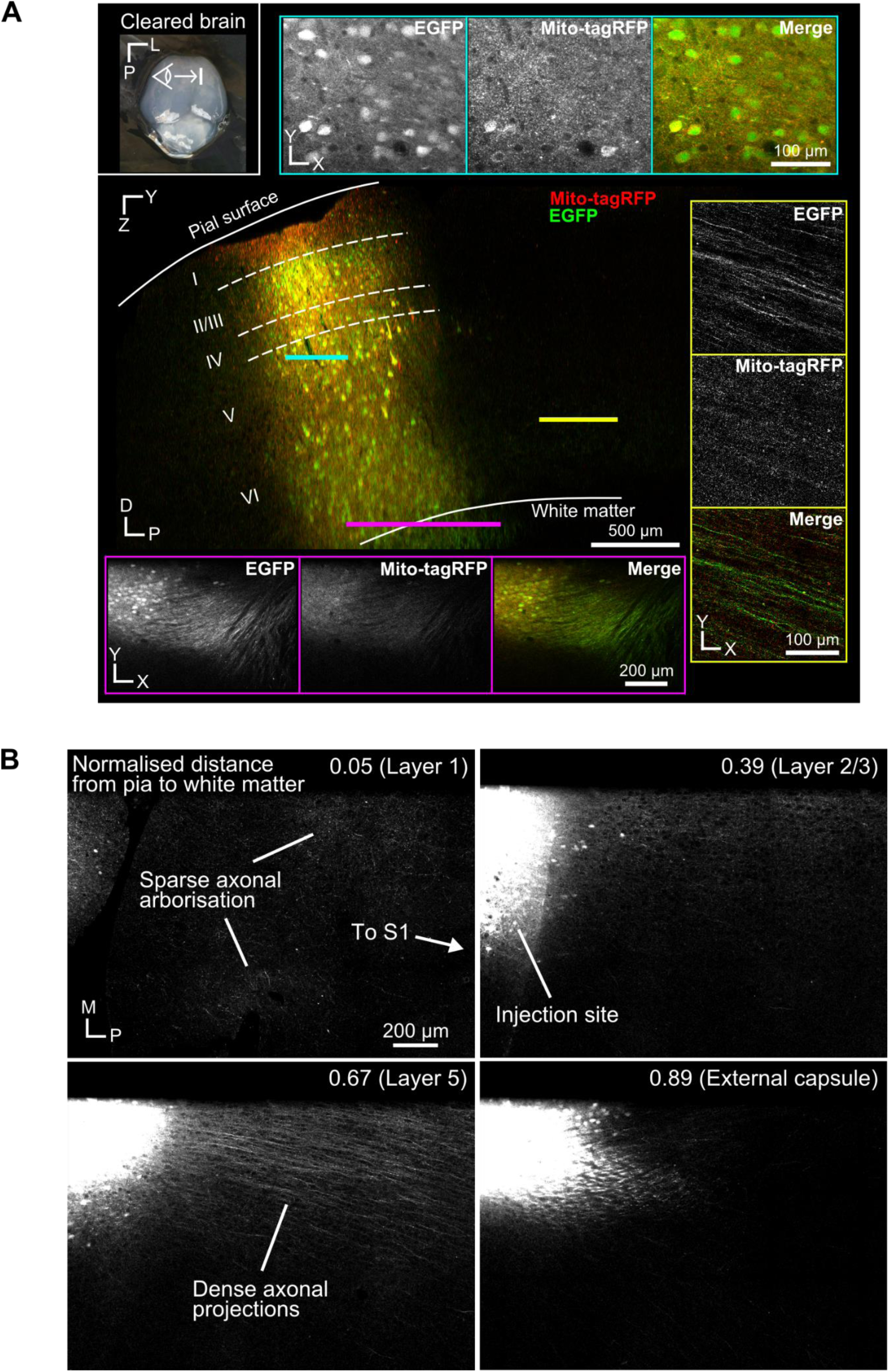
Axonal projections leaving motor cortex. (**A**) (*top-left inset*) low magnification photo of a cleared brain in PBS with the sagittal plane of 2P imaging indicated. (*centre*) 2P image acquired at the viral injection site (sagittal plane). Coloured horizontal lines correspond to inset images with orthogonal imaging planes (horizontal plane). 2P images show the injection site (*top-right inset*), white matter projections (*bottom inset*) and layer V/VI projections (*right inset*). D = dorsal, P = posterior, L = lateral, roman numerals indicate estimated cortical cell layers. (**B**) Horizontal plane through the injection site in cleared brain. (*top-left*) Sparse axonal arborisations in layer 1 of motor cortex. (*top-right*) Over-saturated image showing the injection site at layer 2/3 with sparse axonal arborisation. (*bottom-left*) Dense axonal projections without arborisation emerging from infragranular layers. (*bottom-right*) Axonal projections enter the white matter for subcortical targets. These images indicate that most axonal projections towards somatosensory cortex emerge from layer V/VI. The number in the top-right indicates the normalised distance to the white matter from the pial surface of the cortex. M = medial, P= posterior.

**Figure S3.**
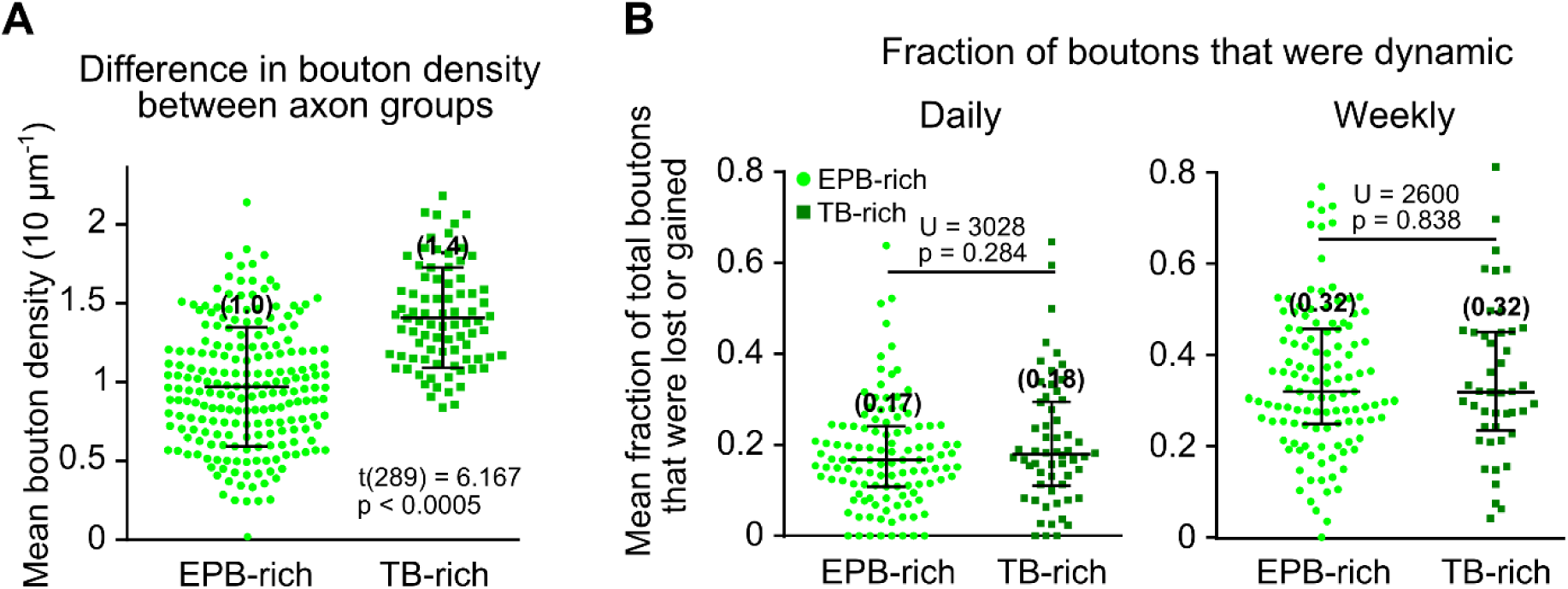
Structural differences along axonal segments were not indicative of bouton turnover. **(A)** Bouton density (mean total density across time) was significantly higher for TB-rich axons compared to EPB-rich axons (mean 1.4 ± 0.3, 1 S.D. and 1.0 ± 0.3 per 10 µm, respectively; independent t-test, p < 0.0005). Error bars ± 1 S.D. Mean indicated in brackets above error bars. (**B**) The fraction of boutons that were dynamic (either lost *or* gained) was not different between EPB-rich and TB-rich axons (mean fraction across daily intervals, 17% and 18%, or weekly intervals, 32% and 32%, respectively; Mann-Whitney U test, p > 0.05). Error bars = inter-quartile range (I.Q.R.). Median indicated in brackets above error bars.

**Figure S4.**
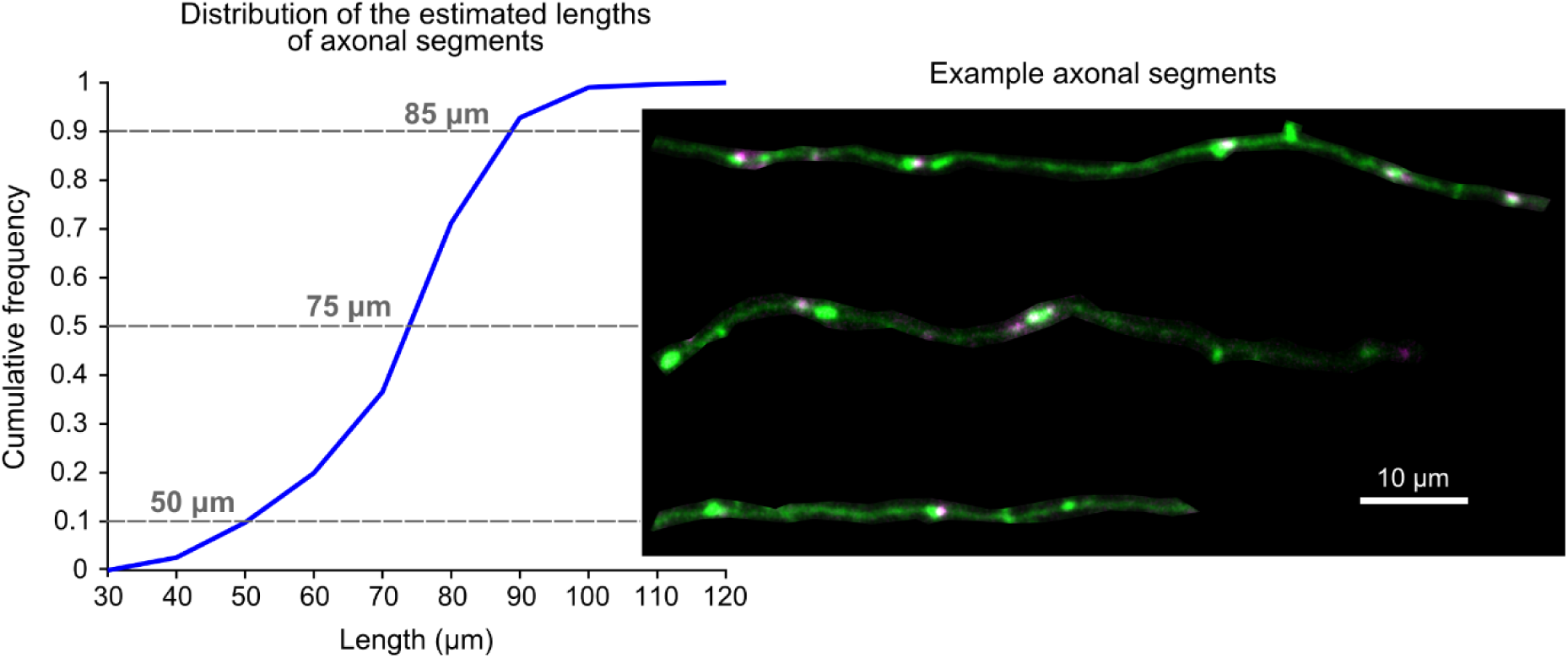
Length distribution of sampled axonal segments. For this study, axonal segments were sampled from 74 x 74 x 30 µm (x, y, z) stacks of 2P images. Each axonal segment length was estimated from a manual tracing made on a 2D projection of each stack (*right*; axons from different fields of view are cropped for presentation). The distributions of estimated axonal segment lengths shows that the median length was 75 µm, with 90% of the sample axonal segments being between 50 and 85 µm (indicated by grey dashed lines).

**Figure S5.**
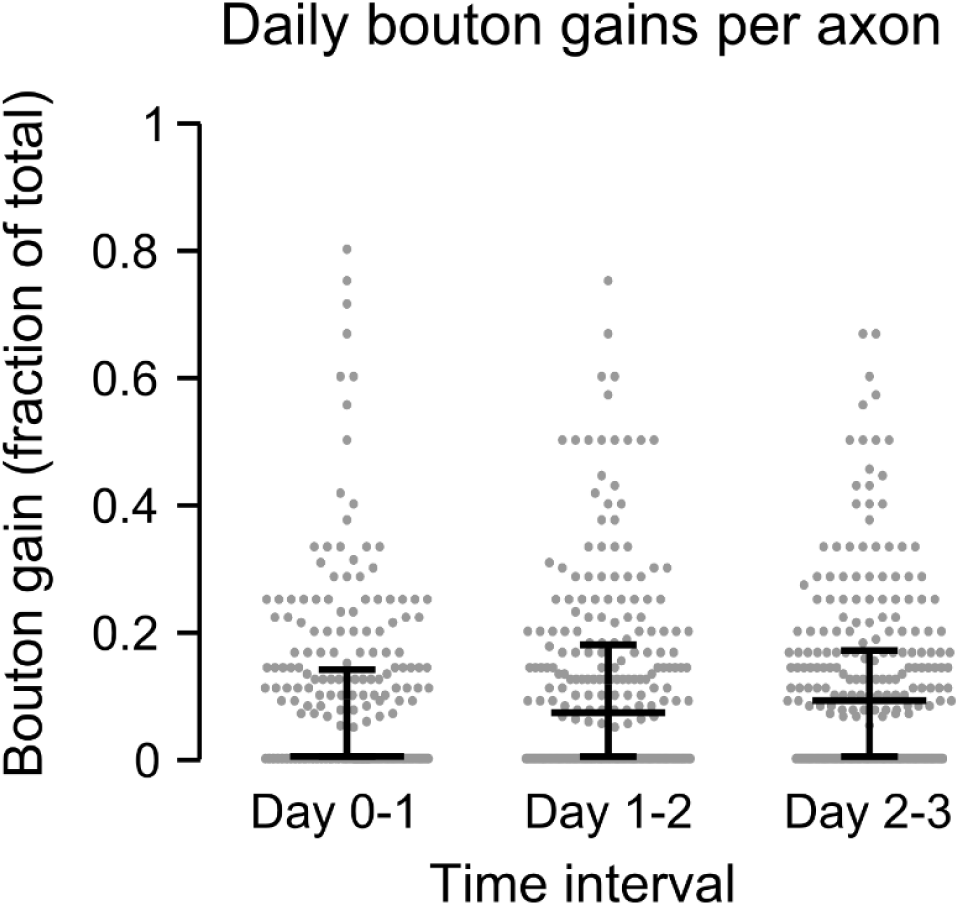
Daily bouton gains as a fraction of the total number of boutons. Over daily intervals the average fraction of boutons gained was 0% (0-14%, I.Q.R.), 8% (0-18%) and 9% (0-17%) (median of the population of axons, n = 249 axons).

**Figure S6.**
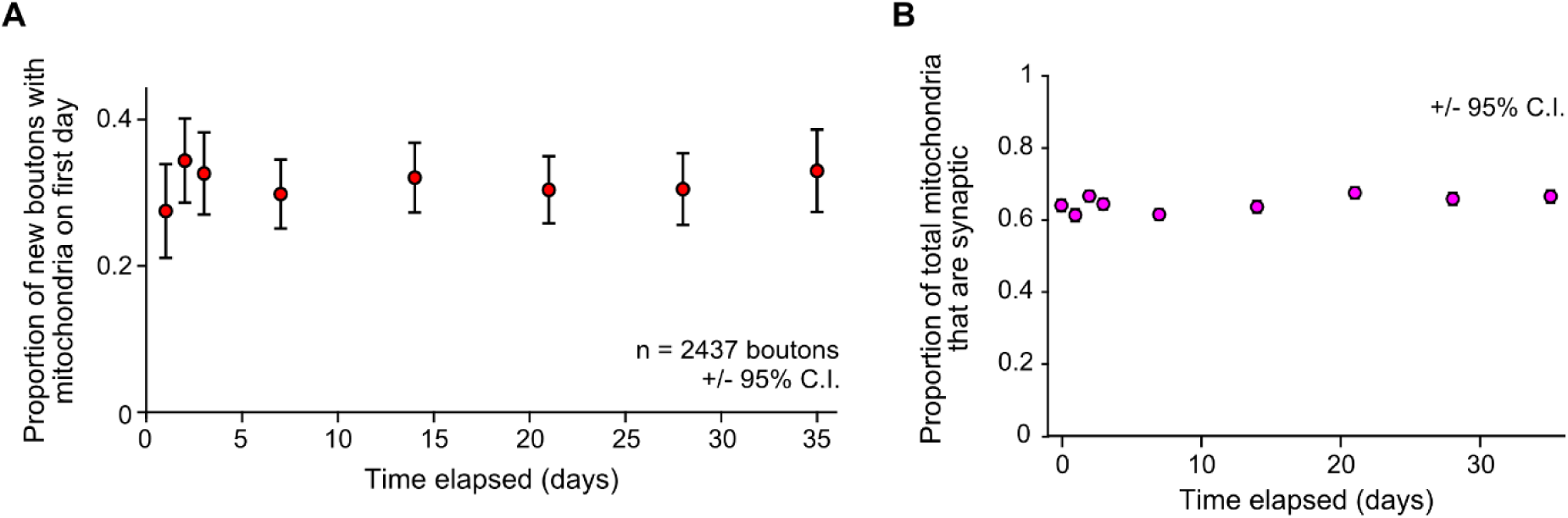
Mitochondrial localisation to presynaptic terminals is consistent for the entire imaging paradigm. (**A**) New boutons formed at every imaging timepoint had the same likelihood of having a mitochondrion nearby, suggesting that increased mitochondrial localisation to long-lived boutons (Figure 4C) is a function of bouton age and not time alone. (**B**) Mitochondria were also no more likely to localise to presynaptic terminals throughout the imaging paradigm.

**Figure S7.**
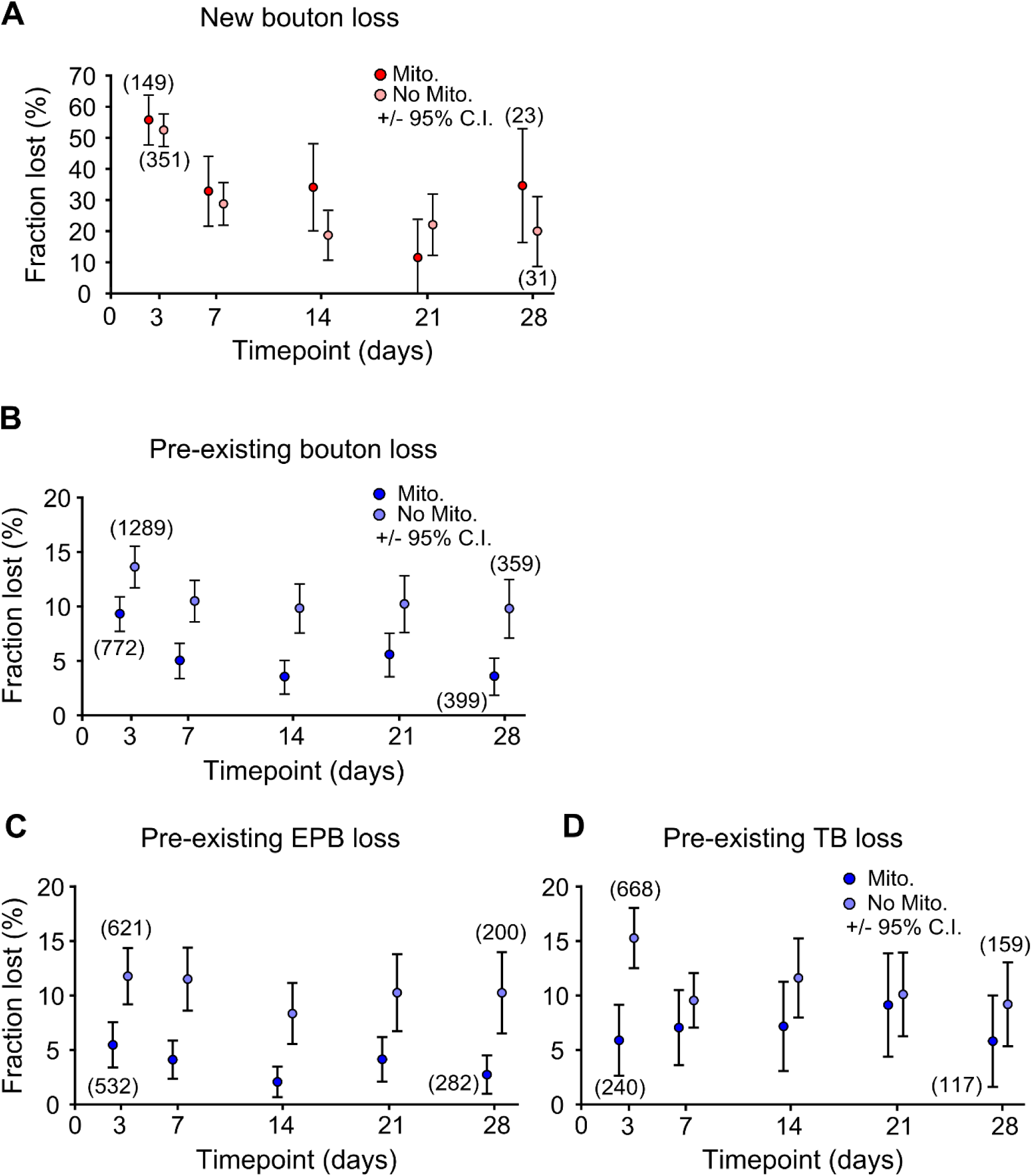
Relationship between mitochondrial presence and bouton loss across time. The number of boutons lost at each timepoint was calculated with relation to the presence of mitochondria, comparisons were made using all the boutons present at each timepoint. (**A**) There was no consistent relationship between mitochondrial presence and bouton loss for newly-formed boutons across time. **(B)** Pre-existing boutons were consistently 30-50% less likely to be lost when mitochondria were localised nearby. (**C-D**) The difference in bouton loss was mainly due to a relationship between mitochondria and EPBs (C), rather than TBs (D). Numbers in brackets indicate the number of boutons in each group (with or without mitochondria) at the start and end of the experiment. The two groups are plotted either side of each imaging timepoint for easier visualisation of error bars.

